# Bat anthropogenic roosting ecology influences taxonomic and geographic predictions of viral hosts

**DOI:** 10.1101/2023.12.12.571362

**Authors:** Briana A Betke, Nicole Gottdenker, Lauren Ancel Meyers, Daniel J Becker

**Author notes:** Correspondence to: Briana Betke.

## Abstract

1. The ability of wildlife to live in anthropogenic structures is widely observed across many animal species. As proximity to humans is an important risk factor for pathogen transmission, anthropogenic roosting may have important consequences for predicting the spillover and spillback of viruses. For bats, the influence of roosting in anthropogenic structures on predicting virus hosting ability and diversity is poorly understood. Such information could be useful for optimizing models of virus outcomes to identify species and locations to target for viral discovery at the human–wildlife interface.
2. We integrate novel roosting ecology data into machine learning models to assess the importance of anthropogenic roosting in predicting viral outcomes across bat species. Additionally, we evaluate how this trait affects the prediction of undetected but likely bat host species of viruses.
3. The importance of anthropogenic roosting varies across viral outcomes, being most important for virus hosting ability and less so for zoonotic virus hosting ability, viral family richness, and zoonotic family richness. Across viral outcomes, anthropogenic roosting is less important than human population density but more important than most family, diet, and foraging traits.
4. While model performance was not affected by inclusion of anthropogenic roosting, models with this novel trait extended the list of undetected host species compared to models excluding this trait. Predicted virus host distributions show distinct spatial patterns between anthropogenic and natural roosting bats, with the greatest proportion of likely novel hosts relative to bat species diversity being anthropogenic roosting bats in Asia.
5. *Synthesis and applications*. These findings suggest anthropogenic roosting has a non-trivial role in predicting viral outcomes in bats, specifically for virus hosting ability. Exclusion of this epidemiologically relevant trait could result in underestimation of predicted hosts, particularly in Asia where hotspots of undetected hosts are mostly able to roost in anthropogenic structures. More broadly, these results demonstrate the importance of evaluating the impact of new data on predicting host–pathogen interactions, even for models that already perform well. Such modeling efforts are particularly relevant for assessing spillover or spillback risks at human–wildlife interfaces.

## Introduction

As emerging infectious disease (EID) outbreaks increase (Meadows et al., 2023; Smith et al., 2014), promptly detecting zoonotic hosts is an important but challenging task. Sampling efforts typically involve resource-intensive field and laboratory efforts to test potential wildlife hosts for the presence of past or active infection with a zoonotic pathogen. Statistical models can minimize logistical and monetary costs by guiding where sampling efforts should be directed at various taxonomic and spatial scales (Becker et al., 2019, 2022; Han et al., 2015; Olival et al., 2017). Further, the resulting predictions can lead to testable hypotheses about mechanisms that drive spillover and spillback events. For example, the prediction of frugivorous bats and primates as likely hosts of ebolaviruses informed further research on ecological and phylogenetic drivers of spillover events, finding evidence from historical data that pteropodid bats may serve as primary hosts and cercopithecid primates as secondary hosts of these viruses in Sub-Saharan Africa (Sundaram et al., 2022, 2024).

The ability to accurately identify potential host species is heavily dependent on the quality and quantity of data used to build each predictive model. Typically, such models use species-level network structure or ecological trait data to classify known hosts from negative or unsampled hosts, often using large, standardized datasets. When new predictor data are provided, it is thus necessary to assess potential changes in model performance to optimize pathogen surveillance. Significant efforts have been made to improve host model performance and predictive ability. Examples include incorporating host competency data (i.e., the ability to transmit a pathogen; Becker & Han, 2021; Mull et al., 2022) to narrow predictions, building models on taxonomic subsets of data to improve predictive capacity (Dallas & Becker, 2021), and assigning pseudo-negatives based on geographic overlap of known hosts to reduce uncertainty and noise in host–pathogen data (Tonelli et al., 2024). While these efforts are strides toward improved model prediction and performance, data deficiencies limit our ability to further improve such models that effectively distinguish hosts from non-host species.

One such data deficiency is information on interfaces where humans and wildlife interact. The observed increase in EID outbreaks has been attributed to greater contact between humans and wildlife where they geographically overlap, known as human–wildlife interfaces (HWIs; Hassell et al., 2017; Jones et al., 2013). Processes such as urbanization increasingly drive some wildlife to live in anthropogenic structures (e.g., buildings, bridges, homes, and tunnels) in response to rapid habitat loss (Adam & Hayes, 2000; Clark, 1994; Imlay et al., 2018; Widdows & Downs, 2016). The propensity of wildlife to live in these anthropogenic structures may have consequences for pathogen transmission by increasing contact between host species and facilitating introduction to novel hosts, including humans. Incorporating data on the occupancy of anthropogenic structures in predictive models could in turn inform sampling efforts by identifying high-priority potential hosts that are most likely to come into contact with humans and locating opportunities for bidirectional pathogen transmission.

Prior meta-analyses and comparative studies have used urban presence and absence data to demonstrate the propensity for urban-attracted species to have greater pathogen diversity (Albery et al., 2022; Gibb et al., 2020; Murray et al., 2019; Werner & Nunn, 2020). However, this body of research relies on broad dichotomies of urban presence or human population density as proxies of human–wildlife contact. These proxies do not adequately represent the physical structures used as HWIs, because they either omit observations of animals in anthropogenic structures (e.g., Jung & Threlfall, 2018) or do not clearly distinguish animals existing in urban areas for long periods of time from those that may only ephemerally visit urban areas (e.g., Albery et al., 2022). Additionally, data documenting visitor and dweller status exists (Santini et al., 2019) but does not comprehensively cover mammals. Such omissions can impact model prediction by identifying host species present in urban areas that have low probability of interacting with humans in ways that facilitate pathogen exposure. For instance, humans can be exposed to pathogens through excess droppings when walking under bridges or in buildings where bats live in high densities (Gerbáčová et al., 2020; Mengistu et al., 2022; Pavlik et al., 2021) as opposed to encountering wildlife that visit urban areas but typically avoid humans such as mountain lions (Suraci et al., 2019). Further, most work focuses on intra-rather than interspecific comparisons, and the coverage of urban presence data at the species-level is poor, challenging our ability to assess synanthropic behavior as a predictor in models. One rare interspecific comparison found that rodents that lived in or near anthropogenic structures increased their probability of being zoonotic hosts, likely owing to factors that relate to their fast life histories (i.e., frequent and large litters, large fluctuations in population size; Ecke et al., 2022). Additional investigation is needed to identify general patterns in the relationship between dwelling in anthropogenic structures and pathogen outcomes across taxa to fully assess risk of both pathogen spillover and spillback events.

Bats and their associated viruses are a timely system to assess the epidemiological importance of wildlife that occupy anthropogenic structures in predictive models. Due to their higher mutation rates and transmissibility than other pathogens, viruses are responsible for most EIDs (Jones et al., 2008). Many bat species play a role in the transmission of these viral pathogens and have well-documented associations with high-profile zoonotic viruses (e.g., Hendra virus, Nipah virus, Marburg virus, SARS-like coronaviruses). For this reason, ample trait data and viral association data exist for bats due to increased surveillance. Previous studies have applied these data to models with the intention of identifying highly predictive traits of viral diversity and detection of novel hosts. This body of work provides a list of known factors such as geographic range size, body mass and size, fecundity, and biogeography (Guy et al., 2020; Han et al., 2016; Plowright et al., 2019) to compare against the importance of anthropogenic roosting.

In addition to data availability, bats are sensitive to environmental changes, and their behavioral ecology, particularly roost selection, is changing alongside urbanization with epidemiologically relevant implications. The redistribution and reduction of available roosting structures due to urbanization have led to four in 10 extant species to occupy anthropogenic structures (Betke et al., 2024). Generally, anthropogenic roosting can affect viral dynamics through increased crowding, fracturing of roosts, and impairment of bat immune responses (Allen et al., 2009; Plowright et al., 2011). For example, Mexican free-tailed bats (*Tadarida brasiliensis*) occupying bridges in Texas showed lower T cell responses than their cave-roosting counterparts (Allen et al., 2009), possibly owing to smaller colony sizes and ectoparasitism in bridges than caves (Allen et al., 2011). Anthropogenic roosting bats are also subject to increased human interactions, resulting in interactions that may further facilitate viral spillover and spillback. For instance, a survey in rural Kenya showed that 38% of humans sharing buildings with bats reported engaging in both direct (i.e., touching, bites, and scratches) and indirect (i.e., urine and feces) contact (Jackson et al., 2024). Furthermore, forced evacuation of bats from their anthropogenic roosts can cause movement into other unoccupied buildings, facilitating spatial spread of viruses such as rabies virus among wild bat populations (Streicker et al., 2013).

Wildlife-occupied anthropogenic structures represent an increasingly critical frontier for bidirectional pathogen transmission between humans and wildlife. Given this potential for direct contact with humans and rapid urbanization occurring globally, we argue that anthropogenic roosting data should be incorporated into models to improve prediction of hosting ability and viral richness. Here, we draw from a novel roosting ecology dataset (natural vs. anthropogenic) compiled in our previous work to compare the importance of anthropogenic roosting to other factors with known associations with viral outcomes (i.e., virus family richness, zoonotic family richness, and hosting ability). We then quantified model performance with and without anthropogenic roosting to test if predictive ability improves with the addition of this epidemiologically relevant and novel trait. Finally, we tested how such models with and without this trait differed in their predictions by also comparing predicted undetected virus and zoonotic virus host species. We expect that inclusion of this would significantly improve hosting and viral richness models due to greater sampling effort of anthropogenic roosting species, more frequent spillover between species (including humans) in these interfaces, and/or impacts on infection dynamics within such species noted above.

## Methods

### Virus data

Virus associations were extracted from The Global Virome in One Network (VIRION) database version 8 (Brookson et al., 2025) and matched to the 1,279 extant bat species from the most recent bat phylogeny, representing 86% of the 1,482 recognized species in the order Chiroptera (Carlson et al., 2022; Simmons & Cirranello., 2023; Upham et al., 2019). VIRION is a comprehensive database that aggregates several host–virus association datasets to create a dynamic and up-to-date collection of host–virus associations with standardized viral taxonomy. These sources include CLOVER, the USAID Emerging Pandemic Threats PREDICT program, and NCBI GenBank. We restricted VIRION to NCBI-verified host and virus species names.

Because serological data may lead to overcounting viral associations given known issues of cross-reactivity (Gilbert et al., 2013), we further restricted data to those virus associations detected by PCR and isolation methods. Two measures of virus richness were derived from VIRION: total viral family richness and zoonotic virus family richness. Total viral family richness is the number of all known viral families for a given bat species (Albery et al., 2022) and zoonotic family richness is the number of viral families for a given bat species found to share at least one virus per family with humans within VIRION. Two binary variables for virus and zoonotic host status were derived from virus species richness to predict potential host species of at least one virus. Any bat species without viral associations in VIRION were designated values of zero (i.e., a pseudoabsense).

### Anthropogenic roosting data

We used an established dataset of anthropogenic roosting ecology manually compiled from existing literature and historical records (Betke et al., 2024). This dataset consists of roosting ecology across 80% (*n* = 1,029) of the 1,279 bat species included in the recent phylogeny and is well represented across bat families and biogeographic regions. Of these 1,029 bat species, 40% roost in anthropogenic structures and 42% exclusively use natural roosts. Roosting in human-made structures is described as a binary classification between species-level use of anthropogenic roosts (e.g., mines, buildings, homes, bridges, tunnels, roofs, and attics) and of exclusively natural roosts (e.g., palm leaves, caves, tree hollows, foliage).

Matching VIRION data to our anthropogenic roosting dataset revealed a wide range of virus richness (0 to 23 virus families). The bat species with the highest virus family richness was *Rhinolophus sinicus*. These data contained 367 bat species with known associations with at least one virus, for which 243 of these species were anthropogenic roosting. Similarly, 222 bat species were found to host at least one zoonotic virus, of which 151 species were anthropogenic roosting. We observed distinct patterns in virus richness across bat families and roosting status (Figure 1A). Descriptive density plots suggested the families Rhinolophidae, Pteropodidae, Mormoopidae, and Miniopteridae had higher identified virus family richness among anthropogenic roosting species; however, these trends were less apparent for zoonotic family richness (Figure 1B).

**Figure 1.**
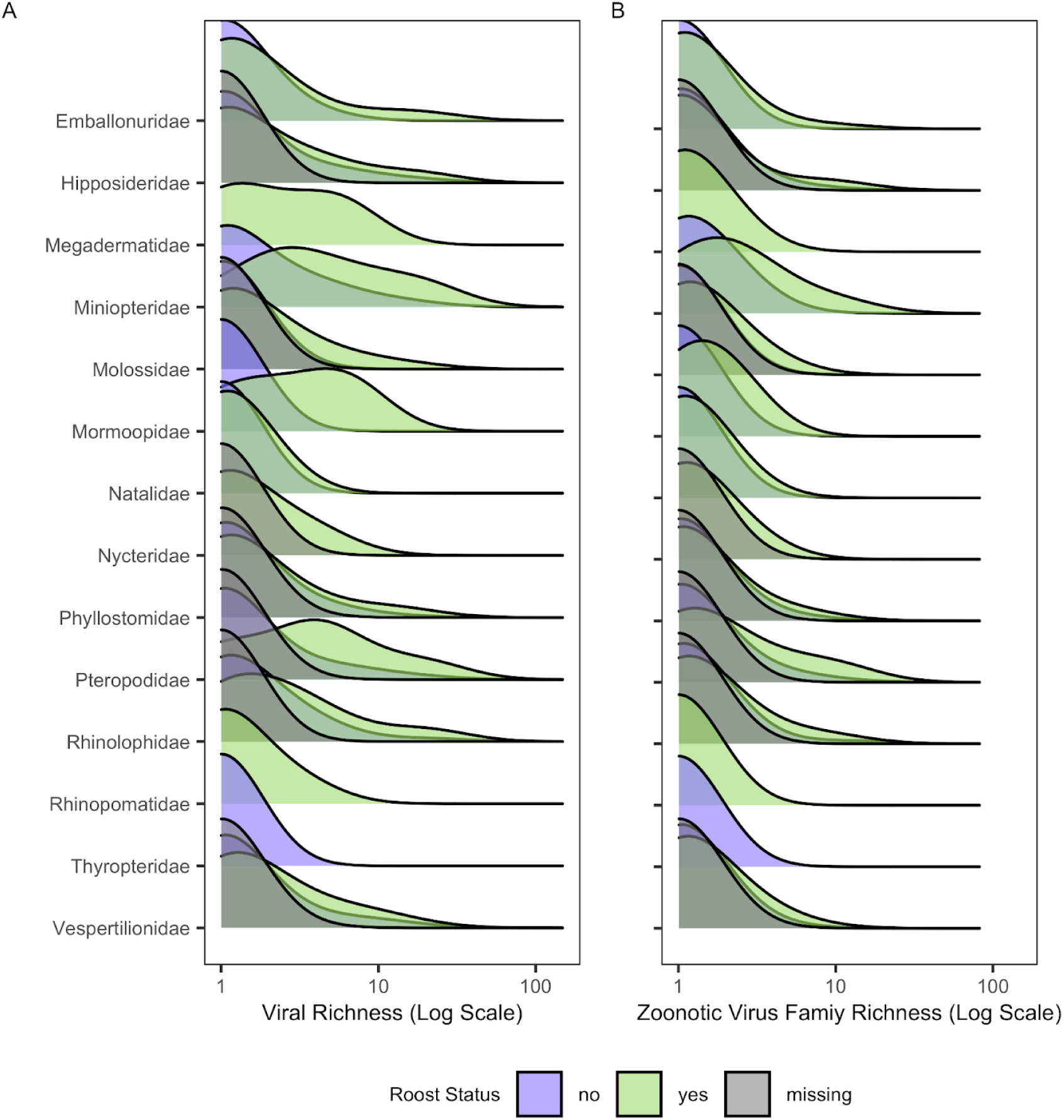
Distribution of anthropogenic roosting and viral outcomes across bat families. (A) Density ridgeline plots of log virus richness of bats faceted by families with observations greater than 5. (B) Density ridgeline plots of zoonotic virus richness faceted by families with observations greater than 5. Distributions are colored by anthropogenic (green), natural (purple), and unknown roosting bats (gray).

### Additional bat trait data

For additional bat traits, including but not limited to those included in prior predictive models of bat virus outcomes (Guy et al., 2020; Han et al., 2016; Plowright et al., 2019; Turmelle & Olival, 2009), we primarily used COMBINE, a recent repository containing 54 mammalian traits aggregated from several commonly used sources (Soria et al., 2021). Traits include important life history variables from PanTHERIA (Jones et al., 2009) and foraging attributes from EltonTraits (Wilman et al., 2014), both of which have commonly been used in prior analyses to identify drivers of pathogen diversity and identify undetected host species (Albery et al., 2022; Becker et al., 2018; Han et al., 2015; Luis et al., 2013; Mull et al., 2022; Olival et al., 2017; Ostfeld et al., 2014). We restricted COMBINE to non-imputed trait data, as imputation of missing trait data may not account for strong biases in coverage (Johnson et al., 2021). Biogeographic areas were expanded to binary variables for species occupancy in each biogeographic region (i.e., Afrotropical, Australian, Indomaylan, Nearctic, Neotropical, Oceanian, and Palearctic).

Additional geographic variables (i.e., geographic area, human population density, mean precipitation, mean monthly actual evapotranspiration rate [AET], mean monthly potential evapotranspiration rate [PET], minimum and maximum latitude, and mean temperature) were collected from the PanTHERIA database. To account for the shared evolutionary history of bat hosts and their impact on viral outcomes, we generated binary variables of all 18 bat families (Han et al., 2016). This trait matrix was merged and taxonomically reconciled with the host–virus association dataset to serve as the trait matrix in this analysis.

### Assessing trait data coverage and sampling effort

We evaluated all trait variables for variance and coverage prior to downstream modeling. Any variables found to have no variation or coverage under 30%, indicating mostly missing values, were removed from the dataset (Figure S1, Supporting Information). We thus excluded 36 out of 99 variables from our analysis, leading to 63 total variables in the dataset in addition to our primary anthropogenic roosting variable (Supporting Information).

Sampling is not equal across hosts and could influence modeling viral outcomes, as some species are studied more due to greater accessibility, proximity to humans, and awareness. Citation count is commonly used as a proxy for sampling bias (i.e., sampling intensity) in trait-based analyses (Albery et al., 2022; Guy et al., 2019, 2020; Mull et al., 2022; Olival et al., 2017). To control for sampling bias in our models, we calculated the number of overall host citations and virus-related host citations for each bat species. A PubMed search was conducted in R with the *easyPubMed* package (Fantini, 2019) to calculate both measures of sampling effort for all bat species in our dataset (Albery et al., 2022; Mull et al., 2022; Olival et al., 2017).

### Boosted regression tree analysis

We built boosted regression tree (BRT) models with the *gbm* package for each of our four response variables to identify traits predictive of overall viral richness, the proportion of zoonotic viruses, and both measures of host status (all viruses and only zoonotic viruses). Bernoulli- or Poisson-distributed error structures were used for binary and count data, respectively. BRTs are machine learning algorithms that determine variables of predictive importance based on an ensemble of regression or classification trees, even in the presence of missing data and non-linear responses, making them ideal for analyzing ecological data (Elith et al., 2008). Trait-based BRTs have also been used to identify unknown hosts of viruses specifically and are found to outperform other modeling approaches for the study of host–virus associations more generally (Becker et al., 2022).

Before fitting BRT models, we partitioned our data between training (80%) and test (20%) datasets using the *resample* package for internal model validation (measured via the area under the receiver operator characteristic curve [AUC] with the *ROCR* package for binary models), across 50 different random data partitions. Models were run with optimal parameters determined via a grid search with 10-fold cross-validation to prevent overfitting (Figure S2, Supporting Information). Resulting variable importance values and performance metrics were averaged across these 50 model runs. We extracted importance rankings for anthropogenic roosting and used a linear model to compare ranks of this variable across our four virus outcomes, using the Benjamini–Hochberg correction to adjust *p*-values for pairwise comparisons (Benjamini & Hochberg, 1995). We used partial dependence plots to visualize the effect of anthropogenic roosting on each viral outcome, after controlling for other features (including citation counts). Additionally, host citation count and virus-related host citation count were modeled with the same explanatory variables as Poisson-distributed responses to determine if viral outcomes are heavily influenced by study effort (Han et al., 2015; Mull et al., 2022; Plowright, et al., 2019).

### Model performance and prediction

Performance metrics were calculated for the appropriate error distributions of each of our four response variables. AUC, sensitivity, and specificity were calculated for our binary models, whereas pseudo *R^2^*was calculated for the richness outcomes. We tested how these performance measures varied between models with and without the anthropogenic roosting variable with unpaired *t*-tests, accompanied by Cohen’s *d* effect sizes and associated *p*-values. The *p*-values were again adjusted with the Benjamini–Hochberg correction. From our models, we predicted viral family richness, zoonotic family richness, and the probability of virus hosting per species after correcting for sampling bias, using mean host citation counts and mean virus-related host citation counts (Becker et al., 2022; Mull et al., 2022). We compared these predictions between models with and without anthropogenic roosting using Spearman rank correlation coefficients and density plots. We then used these predictions to identify unknown host species (i.e., bat species currently without virus associations), where likely overall hosts and zoonotic virus hosts were identified using thresholds that maximize the sum of sensitivity and specificity as determined by the *PresenceAbsence* package (Freeman & Moisen, 2008; Liu et al., 2013). After thresholding, these predicted host species were compared between models with anthropogenic roosting included and excluded. We visualized the spatial distributions of predicted hosts from anthropogenic roosting models if the predictions differed from those of models without anthropogenic roosting. We generated a map of predicted host species using shapefiles obtained from the IUCN Red List database of mammal geographic ranges. To account for bat diversity in our maps, we divided each cell of host species richness by total bat species richness (Guy et al., 2020; IUCN, 2022).

## Results

### Importance of anthropogenic roosting in predicting viral outcomes

Anthropogenic roosting had low-to-medium importance among 62 other features when predicting bat virus outcomes (Figure 2). Our BRTs predicting virus and zoonotic virus host status showed that anthropogenic roosting status is less important than human population density but more important than most family, diet, and foraging traits (see Supporting Information and Figures S3). Roosting ecology was at least as important as human population density change for predicting virus hosting ability and this variable was as important as minimum human population density for predicting zoonotic virus hosting ability (Figure S10). Aligning with past work and subsets of the bat–virus network, other traits relating to geography, life history, and taxonomy were consistently among the most important predictors of virus outcomes (Figure S4, Supporting information).

**Figure 2.**
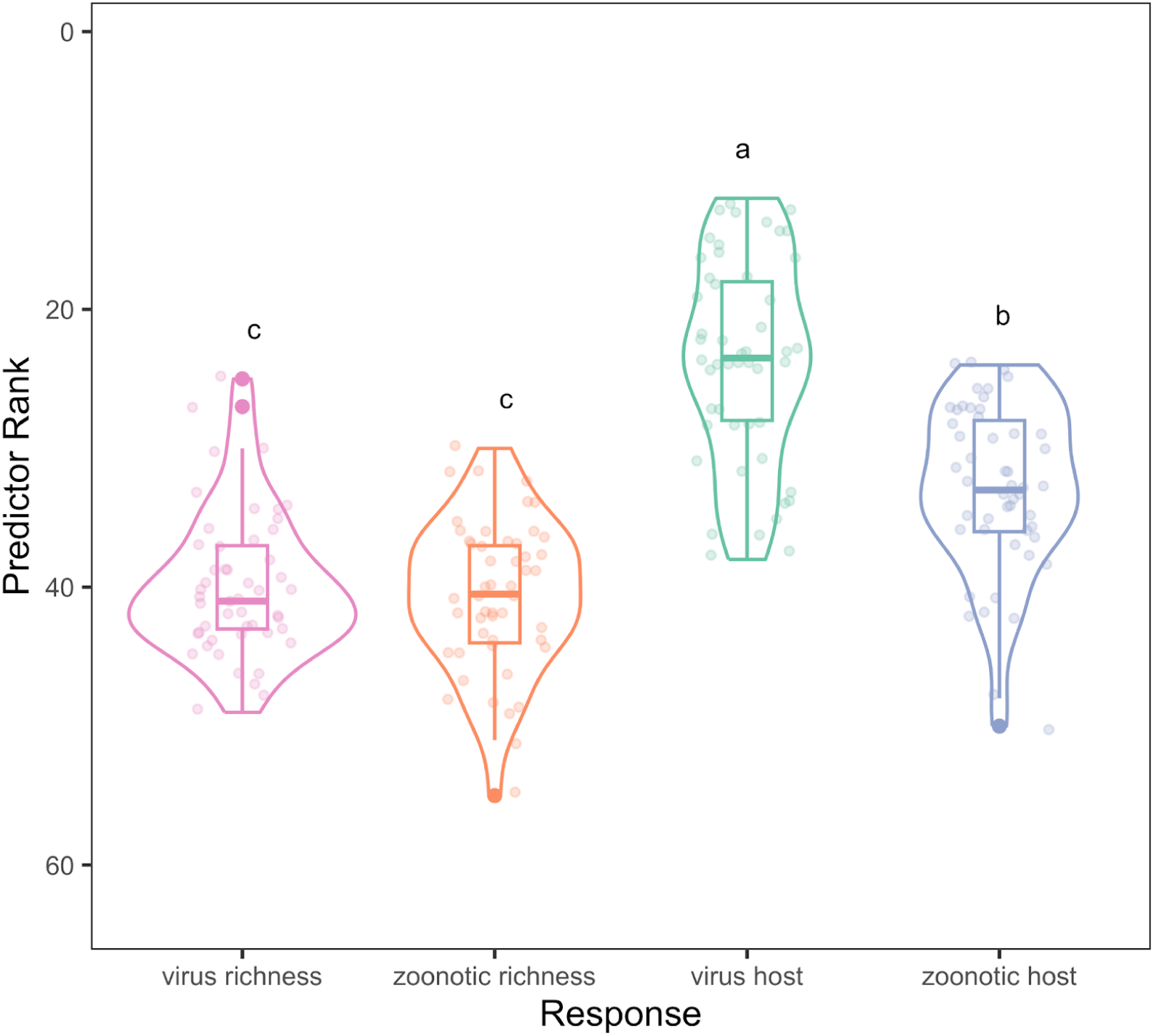
Differences in ranking of our anthropogenic roosting variable across viral outcomes. Colored dots represent rankings for each individual model run (50 random splits of training and test data). Violin plots show the distribution of rankings for each respective viral outcome. Pairwise comparisons between viral outcomes were evaluated with a linear model with posthoc comparison and p-values adjusted for multiple comparisons.

Next, we used variable ranks derived from our BRTs to compare relative importance of anthropogenic roosting across virus outcomes. Anthropogenic roosting ability was ranked highest for predicting virus host status, with a median ranking of 23.5 (range: 12–38) out of 63 variables (Figure 2). The second highest median ranking was zoonotic host status at 33 (range: 24–50). The median rankings for virus family richness and zoonotic family richness models were 41 (range: 25–49) and 40.5 (range: 30–55), respectively. Pairwise comparisons of average ranking for roosting ecology between viral outcomes further confirmed that this variable was more important for predicting viral host status than for predicting zoonotic hosting status, virus family richness, and zoonotic family richness (Figure 2, Table S2). Average ranking of roosting ecology for predicting virus family richness did not differ from zoonotic family richness. Additionally, the average ranking of this variable for predicting zoonotic host status was higher than both family richness outcomes.

Limited relative importance of anthropogenic roosting was also reflected in partial dependence plots for the effect of anthropogenic roosting on virus outcomes while fixing all other trait variables and citation counts. The effect of anthropogenic roosting did not substantially differ from natural roosting across viral richness, proportion of zoonotic viruses, and zoonotic host status (Figure 3). However, anthropogenic roosting had an observable but still weak effect for viral host status, where anthropogenic roosting bats were slightly more likely to host a virus than natural roosting bats (Figure 3).

**Figure 3.**
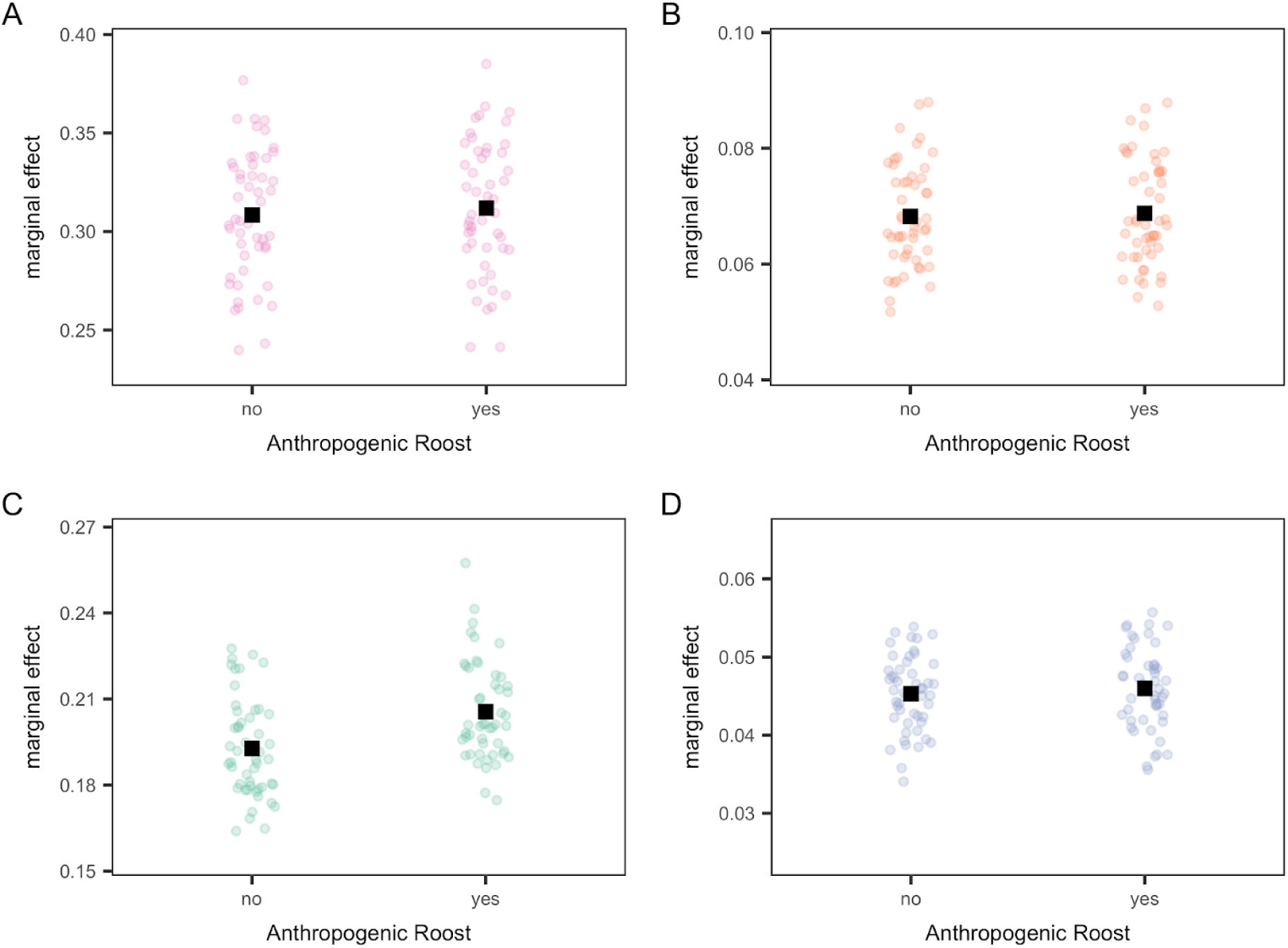
Partial dependence plots for anthropogenic roosting for all viral outcomes. Colored dots represent marginal effects for each individual model run (100 random splits of training and test data), and the means are indicated with black squares. Dot colors correspond to viral outcome, where virus family richness is pink (A), zoonotic family richness is orange (B), virus host status is green (C), and zoonotic host is blue (D).

### Model performance and anthropogenic roosting

BRT models containing anthropogenic roosting predicted overall virus richness with moderately high accuracy (pseudo-R^2^ = 0.52 ± 0.03). Similarly, models with our roosting variable moderately predicted zoonotic family richness (pseudo-R^2^ = 0.52 ± 0.02). Virus hosts were distinguished with high accuracy 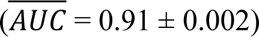, high specificity 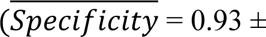 0.333), and moderately high sensitivity 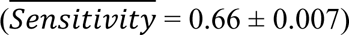 Our models classified zoonotic hosts in a similar fashion to those of virus hosts 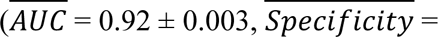 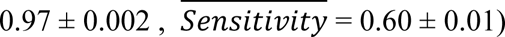.

When comparing the performance of these models to those without anthropogenic roosting, all differences were negligible. Viral richness models without anthropogenic roosting had equivalent performance (pseudo-R^2^ = 0.53 ± 0.03; *t* = -0.3 *p* = 0.97; Cohen’s *d* = -0.06). We observed this lack of difference in performance for zoonotic family richness models (pseudo-R^2^ = 0.53 ± 0.02; *t* = -0.14, *p* = 0.97; Cohen’s *d* = -0.03) and zoonotic host status 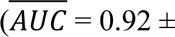 0.003; *t* = 0.18, *p* = 0.97; Cohen’s *d* = 0.04). Similarly, sensitivity 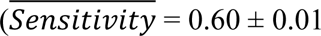; *t* 0.05; *p* = 0.9; Cohen’s *d* = 0.01). and specificity (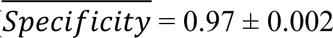; *t* = -0.04, *p* = 0.97; Cohen’s *d* = -0.007) did not differ between models for zoonotic host status. Virus host models without anthropogenic roosting also did not vary in performance (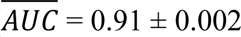; *t* = 0.23, *p* = 0.97; Cohen’s *d* = 0.05), specificity (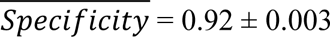; *t* = 0.51, *p* = 0.97; Cohen’s *d* = 0.10), or sensitivity (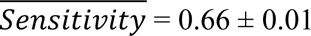; *t* = -0.08, *p* = 0.97; Cohen’s *d* = -0.02).

To assess if the trait profiles of viral outcomes were driven by sampling effort, we ran BRT models predicting both our citation count variables using the same trait predictors as all other models. We found that the included traits poorly predicted both host citation count (pseudo-R^2^ = 0.13 ± 0.07) and virus-related host citation count (pseudo-R^2^ = 0.02 ± 0.09).

### Anthropogenic roosting and model predictions

Predicted virus family richness, zoonotic family richness, virus hosting status, and zoonotic virus hosting status did not vary between models with and without anthropogenic roosting (ρ = 1, *p* < 0.001 for all responses; Figure S9). Despite predictions being very similar between models with and without anthropogenic roosting, the density of model predictions varied by roosting ecology for each viral outcome. We observed greater predicted values for anthropogenic roosting species for both non-binary models (i.e., virus family richness and zoonotic family richness; Figure 4) and binary models (i.e., viral hosts and zoonotic hosts; Figure 5).

**Figure 4.**
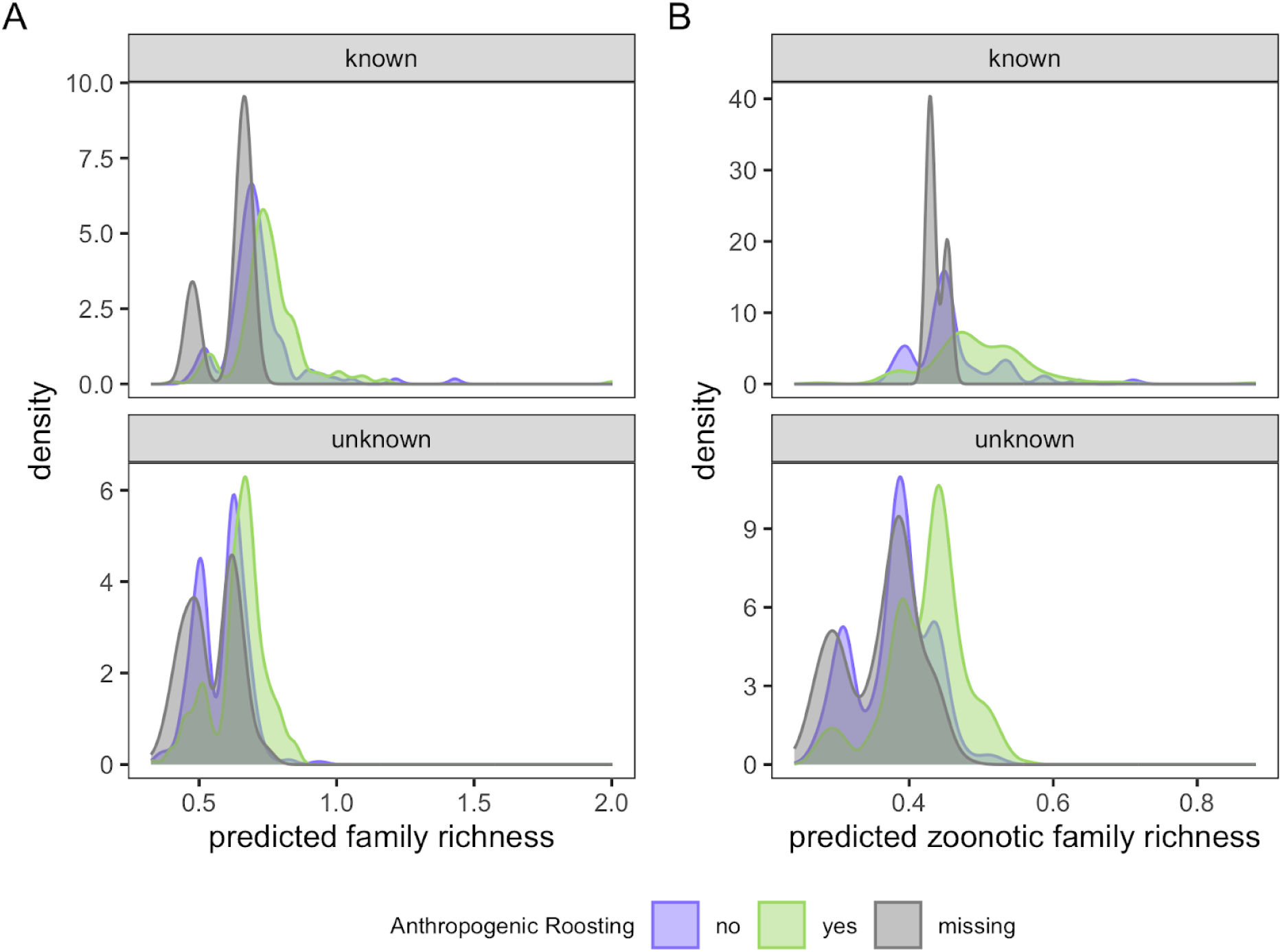
Densities of predicted viral family richness of viruses for models with anthropogenic roosting as a variable. The predicted values for known and unknown bat species from virus family richness models (A) and zoonotic family richness models (B), stratified by roosting status. Natural roosting, anthropogenic roosting, and unknown roost status hosts are indicated by purple, green, and grey, respectively. The x-axes use cubed root transformation for both virus family richness and zoonotic family richness models.

**Figure 5.**
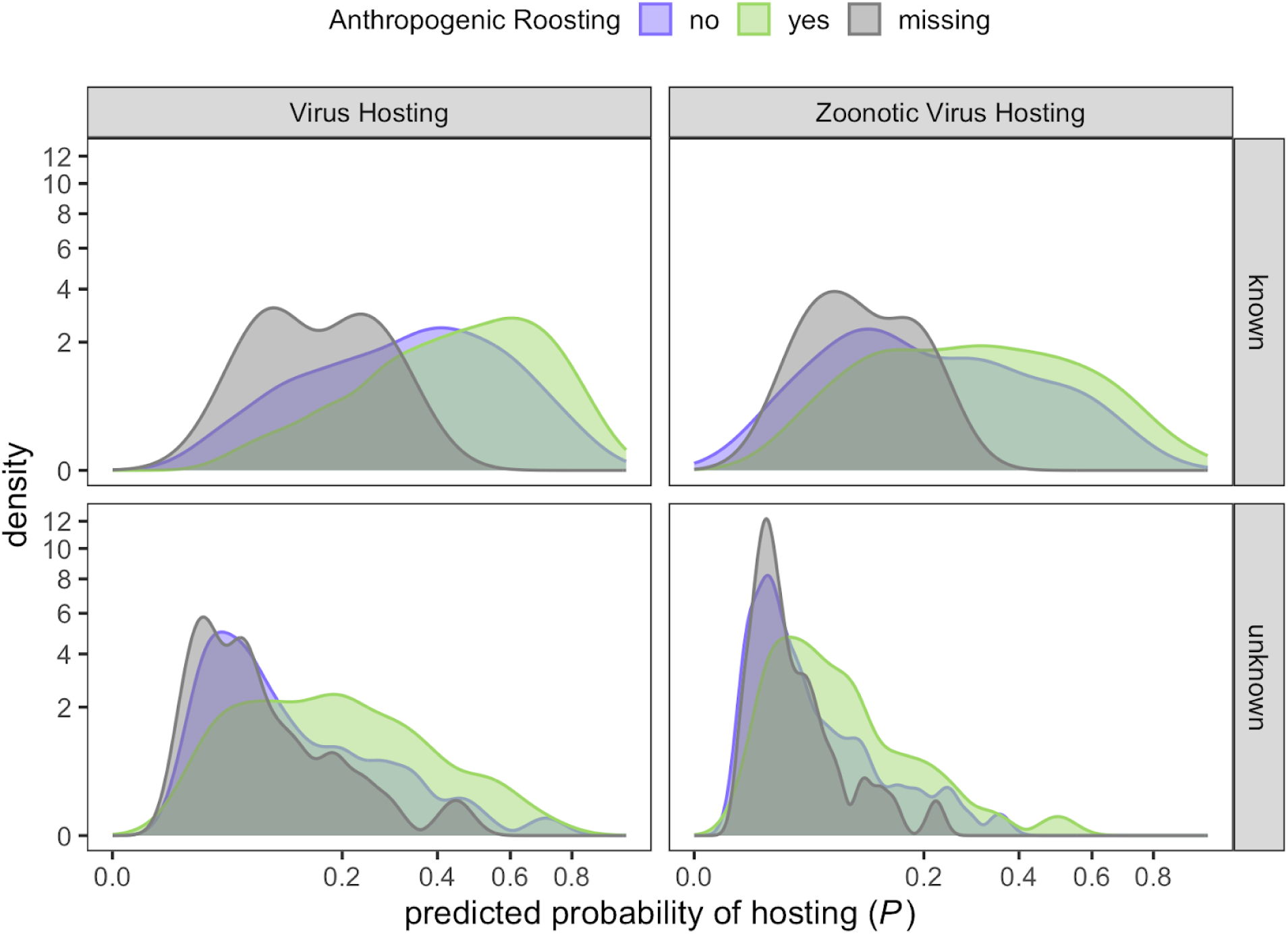
Predicted probability of hosting a virus for models with anthropogenic roosting as a variable. The predicted probabilities of hosting of known and unknown hosts from virus host and zoonotic virus host models, stratified by roosting status. Natural roosting, anthropogenic roosting, and unknown roost status hosts are indicated by purple, green, and grey, respectively. All axes are plotted on a square root scale.

We further compared the predictions of host models with and without anthropogenic roosting. Virus host models with roosting ecology as a predictor identified 134 undetected hosts, whereas models without this trait predicted only 128 undetected hosts (Table S3). Both models were in agreement on 127 host species. Seven of the 134 species were not predicted by models without this trait, all of which were anthropogenic roosting. There was one species uniquely predicted by models without anthropogenic roosting and the roosting status of that species is unknown. Our zoonotic host models with roosting ecology predicted host 150 species in common with those predicted by models without roosting ecology. However, there were six host species not shared between the models. Models without roosting ecology identified four host species not predicted by models with roosting ecology, and the remaining two were only predicted by models with this trait.

When we stratified predictions from host models with roosting ecology by bat family, the Vespertilionidae contained the highest proportion of novel virus and zoonotic virus hosts (52/134 and 59/152 species, respectively). The Phyllostomidae held the second highest number (40/189 overall virus host species and 26/152 zoonotic host species). Most undetected virus hosts were predicted to be in the Neotropics and Afrotropics, with 42 and 40 species out of 134 species, respectively. The Neotropics and Afrotropics also contained the most undetected zoonotic hosts, with 49 and 43 of the 152 predicted species, respectively. When we stratify predictions by anthropogenic roosting status, 83 of the 134 host species identified by host models with roosting ecology were anthropogenic roosting. Of the 152 zoonotic host species predicted by models with roosting ecology, 95 were anthropogenic roosting (Table S4).

We lastly mapped the distribution of known and novel hosts predicted from the virus host model scaled by total bat diversity to observe spatial differences in the distribution of anthropogenic and natural roosting hosts (Figure S11). Novel anthropogenic roosting hosts largely had distinct distributions from the novel hosts that exclusively roost in natural structures. This result was most apparent in the Arabian Peninsula, North American deserts, and perimeter of the Tibetan Plateau, where hotspots of predicted anthropogenic roosting virus hosts are located. When mapping predicted hosts from each model according to their anthropogenic roosting ability, we found that excluding anthropogenic roosting underestimates the number of host species in Asia (Figure 6).

**Figure 6.**
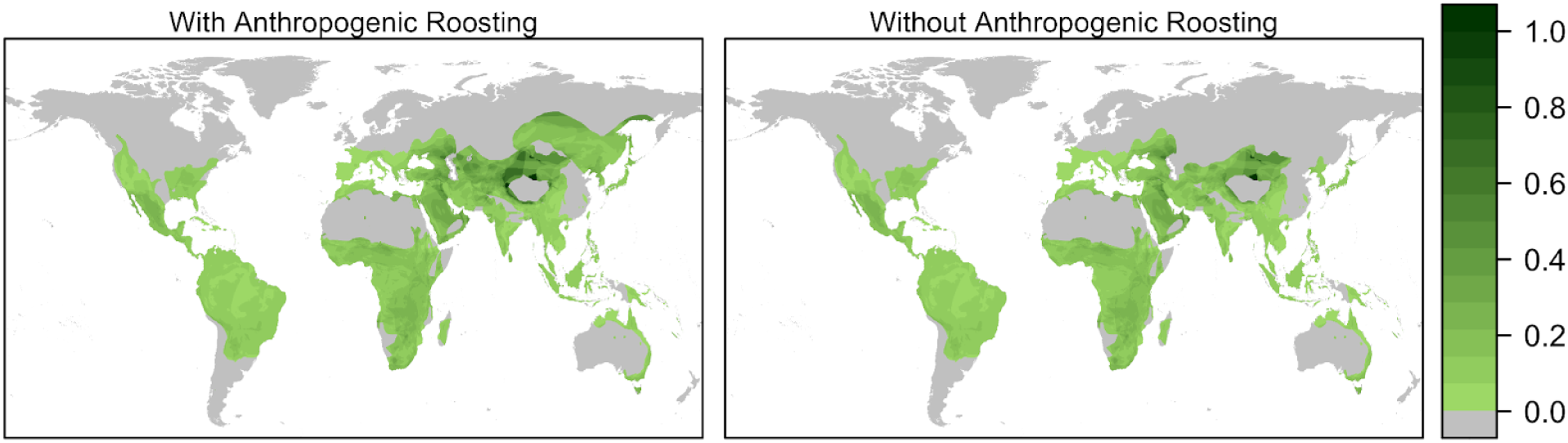
Distribution of predicted virus hosts for anthropogenic roosting species. Left Panel: A map of the anthropogenic species predicted to be hosts for virus hosting models including anthropogenic roosting as a variable. Right Panel: A map of the anthropogenic species predicted to be hosts for virus hosting models without anthropogenic roosting as a variable. Both plots have been scaled by total bat diversity. Darker green regions show concentration of predicted host species when accounting for bat diversity.

## Discussion

As the rate of pathogen emergence increases (Meadows et al., 2023; Smith et al., 2014), early detection of hosts is an important first step towards better anticipating spillover events.

Predictive models using host traits assist with these efforts by providing targeted guidance for surveillance (Becker et al., 2022; Plowright et al., 2019). Such models rely on large, standardized trait datasets and should be evaluated when new information becomes available to ensure optimal predictive ability. In this study, we assessed model behavior with the addition of novel predictor data on the interfaces where humans and wildlife interact. Rather than a broad urban-dwelling criterion (Albery et al., 2022; Santini et al., 2019) or variables such as human population density within a species’ range to approximate human–wildlife contact (Han et al., 2015, 2016; Olival et al., 2017), we focus on the ability of a species to live within human-made structures to directly assess relationships at the human–wildlife interface. Bat roosting ecology, particularly the structures in which bats roost, likely serve as important points of cross-species transmission, given bats’ involvement in the several emerging infectious disease events (Lunn et al., 2024; McKee et al., 2019). Overall, our results suggest that this aspect of bat roosting ecology is only moderately important in predicting viral outcomes (i.e., virus family richness, zoonotic virus family richness, host status, and zoonotic host status) but that the inclusion of this trait provides different taxonomic and geographic predictions of virus hosts compared to models without this trait.

The ability of bat species to use anthropogenic roosts was not the most important nor the least important predictive variable across viral outcomes, especially for predicting undetected hosts of at least one virus. Prior models using different facets of roosting ecology similarly found that roost type did not rank highly in predicting virus diversity (Guy et al., 2020). Although we cannot directly compare our findings to prior work due to differences in roost type categorizations and data coverage, temporal shifts in viral diversity records (Gibb et al., 2022), and modeling strategy, these results may signal a wider trend in the role roosting ecology plays in predicting species-level trends in bat viral diversity. Further inspecting the influence of our roosting ecology variable on predicting viral outcomes showed that anthropogenic roosting bats had a slightly increased probability of hosting at least one virus. Taken with the results of other viral outcomes, our findings suggest that anthropogenic roosting bats have slightly greater probabilities of hosting viruses than natural roosting bats but do not differ in their viral richness. This result deviates from prior work showing that urban-adapted mammal species host greater viral richness (Albery et al., 2022). Our variable importance and marginal effects plots did not strongly support anthropogenic roosting as a major driver of zoonotic outcomes in particular, suggesting that anthropogenic dwelling may not play as critical of a role in predicting zoonotic hosts for bats (as compared to rodents; Ecke et al., 2022).

The overall weak patterns in variable importance and marginal effects observed here may be explained by trait profile differences in bat species with high viral family richness and positive host status from that of anthropogenic roosting bats. In a previous study (Betke et al., 2024), anthropogenic roosting ability was best described by a combination of geographic, climatic, reproductive, and morphological traits. Notably, the top five predictors of this roosting trait were geographic area, habitat breadth, two climatic variables (i.e., mean precipitation and AET), and litter size. Variable importance plots show that the variables most predictive of viral hosting and zoonotic hosting align more with that of anthropogenic roosting ability than those that predict viral family richness (Figure S12). With the exception of citation count, host models show that geographic area is the most important variable for determining virus hosting status, whereas human population density is the most important variable for predicting both virus family and zoonotic virus family richness. In line with these results, human population density did not rank among the top 10 variables for describing anthropogenic roosting ability. Additionally, important climate variables (i.e., AET and PET) found to describe anthropogenic roosting bats also had higher variable importance for viral hosting models than for richness models. The small difference in marginal effects between anthropogenic and natural roosting bats for viral hosting may thus reflect this stronger alignment with anthropogenic roosting traits. Habitat breadth and other dietary variables (i.e., percentage of plants and fruit in diet) were among the most important variables describing anthropogenic roosting but were not highly predictive of any virus outcome. These differences in variable importance suggest that while anthropogenic roosting bats possess many traits that are associated with greater hosting status, these traits are not unique to anthropogenic roosting.

The inclusion of roosting ecology did not substantially improve model performance across viral outcomes, possibly owing to similar models predicting bat virus outcomes already performing well prior to the inclusion of this trait. We built upon similar methodologies from other models of the viral diversity and host status of bats that also show high predictive ability (Becker et al., 2022; Guy et al., 2020; Han et al., 2016; Plowright et al., 2019). This could additionally explain how predictions of viral outcomes from our models incorporating roosting ecology did not substantially deviate from our models without this trait, with the exception of predictions of novel hosts in Asia. However, predictions from virus host models with this trait extended the list of undetected hosts, despite the small improvement in performance.

Differences in predicted hosts across virus and zoonotic host models reveal that roosting ecology had a greater impact on broader virus host prediction than zoonotic host prediction. Virus host models with roosting ecology predicted seven undiscovered hosts than the analogous model without this trait, whereas zoonotic host models with roosting ecology only detected two species not found by analogous models without this trait. This increase in undetected hosts, all of which were anthropogenic roosting, suggests that exclusion of roosting ecology in predictive models could lead to false negatives in terms of detecting hosts of at least one virus. Accurately identifying likely virus hosts is especially important both for study design, given the financial and logistical costs of sampling (Plowright et al., 2019), and for bat conservation, by minimizing unnecessary sampling of vulnerable species. For the latter point, anthropogenic roosting bats may be especially vulnerable to biodiversity loss and are further threatened by forced evacuation due to their reliance on human-made structures (Betke et al., 2024). Fourteen of the 134 undetected virus hosts predicted by models with roosting ecology had near threatened or threatened (i.e., vulnerable, endangered, or critically endangered) conservation status, with 10 of these 14 being anthropogenic roosting species. Bat species with anthropogenic roosting ability also represent the majority of predicted hosts with decreasing population trends. Seventy-eight percent of species with declining population trends have anthropogenic roosting ability (14/19). These results emphasize the need to develop virus surveillance efforts with bat conservation in mind.

Most undetected viral hosts were located in the Neotropics and Afrotropics (42 and 40 of the 134 predicted hosts, respectively). However, this disproportionate number of hosts is also a function of greater bat diversity in these regions, such that we did not identify hotspots in the Neotropics or Afrotropics relative to global patterns in bat species richness (Figure S11). Previous work has shown similar patterns for the Neotropics when accounting for bat richness (Guy et al., 2020). In contrast to previous work, northern Africa did not contain hotspots of novel hosts after scaling for bat diversity in our analysis (Figure S11). Novel hosts, particularly those that are anthropogenic roosting, are disproportionately concentrated in the areas surrounding the Tibetan Plateau, located in the Indomalayan and Palearctic realms. These hotspots have not been identified in previous studies. Sampling bats in these hotspots, identified by including roosting ecology data, could serve as a useful starting point to geographically expand sampling effort while also prioritizing areas where bat hosts represent a greater proportion of bat diversity.

While we show that including anthropogenic roosting into models of bat–virus interactions can affect taxonomic and geographic foci of risk, inferences from our analysis about the ability for undetected hosts to transmit viruses are limited by the kinds of viral data available. Due to a deficiency in the availability of virus isolation data, we do not distinguish between PCR- and isolation-based detections when calculating viral outcomes from the VIRION dataset. Defining host–virus associations more strictly, such as those based on host competence, and empirically testing predicted hosts for viral competency, could enhance model predictions and advance our understanding of spillover and spillback risks (Becker et al., 2020; Mull et al., 2022). Furthermore, using models informed by host competence may help mitigate stigma against bats labeled as reservoir hosts solely based on serological or PCR evidence (Shapiro et al., 2021; Weber et al., 2023).

This study serves as a starting point for understanding the impact of anthropogenic roosting on cross-species transmission and provides several directions for future work. To evaluate anthropogenic roosting at the host species scale, we collapsed the bat–virus network, leading to a loss of information relating to viral traits. The inclusion of viral characteristics such as nucleic acid (DNA vs RNA virus), enveloped vs non-enveloped virus, or transmission mode (i.e., vector, environmental, direct contact) could affect the importance of anthropogenic roosting for predicting associations with particular viruses. Future models could focus explicitly on link prediction to better integrate both host and virus characteristics into analyses of the host–virus network (Albery et al., 2021; Dallas et al., 2017). Additionally, our modeling approach with roosting ecology and extensions into link prediction could readily be adapted to analyses of other pathogens. Bat–bacterial associations are especially understudied, as research effort is increasingly focused on determinants of viral outcomes, but can also have important implications for human and bat health (Shaw et al., 2020; Szentivanyi et al., 2023). Such analyses could further guide focused field studies to both test pathogen hosting predictions and to understand how anthropogenic roosting affects viral dynamics.

In conclusion, we leveraged recently compiled, novel data on roosting ecology to explore how anthropogenic roosting influences viral outcomes across bat species. While roosting ecology was not a highly predictive trait, our results show that predicted viral hosts are distinctly distributed based on their roosting ecology. Our findings also support anthropogenic roosts as a meaningful interface where bat–human contact (and therefore cross-species transmission) occurs (Jackson et al., 2024; Streicker et al., 2013). A better understanding of how anthropogenic roosting influences bat–virus interactions through the application of macroecological tools and further integration with studies of bat–human interactions within ecological, cultural, behavioral and socioeconomic contexts can provide holistic frameworks to understand as well as prevent transmission of bat-borne viruses and guide field studies in ways that minimize invasive sampling of vulnerable wildlife species.

## Author Contributions

Conceptualization, D.B., B.B.; Methodology, D.B. and B.B.; Formal Analysis, B.B.; Investigation, B.B. and D.B.; Data Curation, B.B. and D.B.; Writing–Original Draft, B.B. and D.B., Writing –Review & Editing; B.B., D.B., L.M. and N.G.; Supervision, D.B.

## Acknowledgements

B.B. and D.J.B. were supported by funding to the Viral Emergence Research Initiative (VERENA) Institute, including the National Science Foundation (DBI 2515340). B.B. was additionally supported through the National Science Foundation Postdoctoral Fellowship in Biology (DBI 2305782). D.J.B was additionally supported by the National Institutes of Health (R01AI185127, R03AI188200).

## Data Availability

The viral association dataset used in this analysis can be accessed at the VIRION GitHub repository (https://github.com/viralemergence/virion). All code and data associated with this analysis can be found at the bathaus GitHub repository (https://github.com/viralemergence/bathaus)(Betke et al., 2023).

