## Supplementary Information for "Bat anthropogenic roosting ecology influences taxonomic and geographic predictions of viral hosts"

**Optimizing Boosted Regression Tree Models**

A grid search of parameters was conducted for all response variables with full trait data. BRTs were fit for each combination using the *gbm* package (Greenwell et al. 2020). For all models, we selected three interaction depths (2,3,4) and three learning rates (0.01, 0.001, 0.0005) across a selection of trees dependent on the type of BRT model (regression or classification tree). For the regression tree models (i.e., virus family richness and zoonotic virus family richness), we selected 5000 and 15000 as initial trees. The classification models were tested with the same initial trees as the regression models with the addition of 25,000. Combinations where there was a low number of trees (5000) and small shrinkage were removed (0.001, 0.0005) as smaller shrinkage values typically require a larger number of trees. Each combination had up to 10 unique test/training splits to account for variation in data splits. A total of 90 combinations were run for the regression models and 120 for the classification models. We calculated RMSE to evaluate performance of regression tree models. For classification trees, we used AUC, sensitivity, specificity. AUC values were calculated with the *ROCR* package (Sing et al. 2005). Sensitivity and specificity were calculated using the *InformationValue* package (Prabhakaran 2016). To select the optimal parameters, we calculated the median/average for all performance metrics across seeds for each combination of parameters. For both richness models, an interaction depth of 4 decreased error (Figure S2). RMSE was lowest for both richness outcomes with an initial 5000 trees, an interaction depth of 4, and a learning rate of 0.01. Parameters for the binary models appeared to perform similarly across interaction depths (Figure S2), thus the selection of parameters came down to the minimization of computational time while retaining optimal performance. Virus hosting models showed the number of 15,000 trees, interaction depth of 4, and learning rate of 0.001 to be optimal based on AUC. However, the combination of 5,000 trees, interaction depth of 2 and learning rate of 0.01 were selected due to maximizing specificity at the cost of slightly lower AUC while reducing computational time and no cost to sensitivity. For the zoonotic hosting models, 5,000 initial trees, interaction depth of 2, and shrinkage of 0.001 were found to be optimal parameters. Similar to the virus reservoir models, this selection was based on slightly sacrificing AUC for higher specificity and sensitivity. No training and testing splits reached the 5,000 maximum number of trees for any outcome, the max number of trees were reduced to 2,500 to further reduce computational time.

**Top ecological predictors**

Across all outcomes, citation count and virus citation count were the top two most important variables, with increasing marginal effect as citation count increased (Fig. S3-4). Bat species with high virus richness exist at low to medium human population densities, are located in areas with moderate water loss, and are distributed across longitudes except in the extremes of the eastern hemisphere (Fig. S5). In contrast, species with high zoonotic family richness are located in areas with low to moderate human population densities, can withstand higher upper elevation limits, and exist across almost all minimum latitudes (Fig. S6). Virus hosts tend to have large geographic ranges, exist in areas with moderate water loss, and have maximum latitudes in the northern hemisphere (Fig. S7). Zoonotic virus hosts are similarly defined to virus hosts geographically, with the exception of larger body lengths for zoonotic hosts and larger forearm lengths for overall virus hosting (Fig. S8).

### Figures

**Figure S1. Coverage of traits considered for inclusion for this analysis (n = 83).** The plot shows the frequency of trait variables by percentage of coverage across bat species. The dotted line is the 30% cut off threshold for inclusion in our trait matrix.

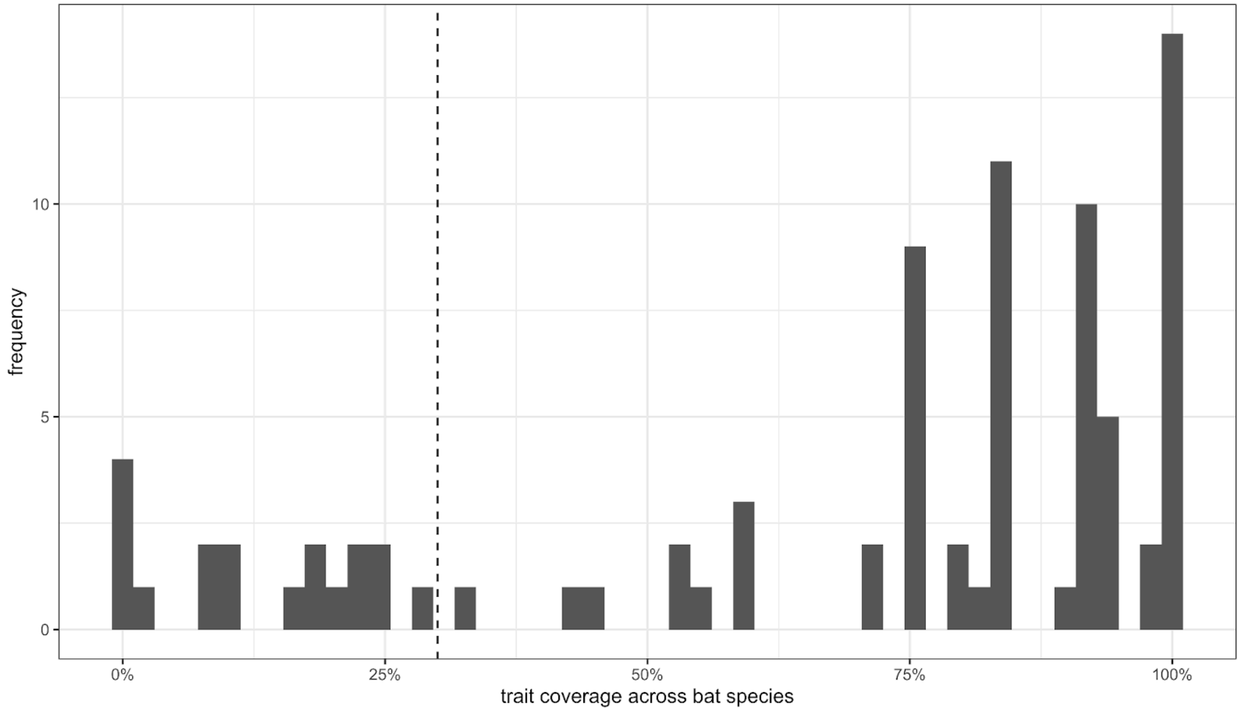

**Figure S2. Performance of tuning parameters in BRTs via grid search**. Test RMSE plotted for poisson and gaussian distributed error models. For binary reservoir status models, test AUC, sensitivity, and specificity were plotted. Learning rates are shown on the x-axis and colors correspond to interaction depths.

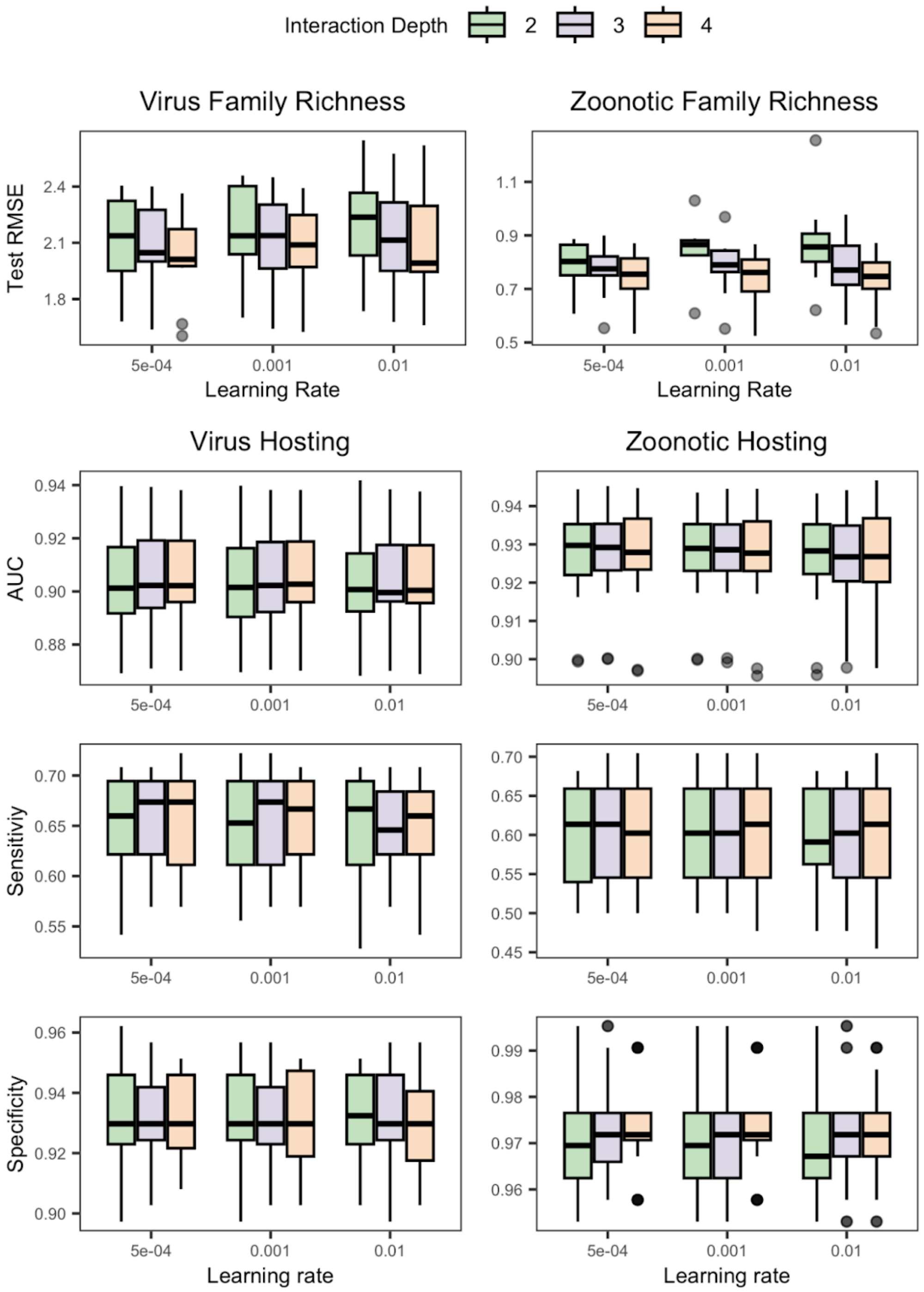

**Figure S3. Variable influence (square root scale) for predicting virus family richness and zoonotic virus family richness.** Bar height shows the importance of each predictor averaged across 100 unique test data splits. Error bars indicate the variation across models. Axes of relative importance are plotted on square root scale.

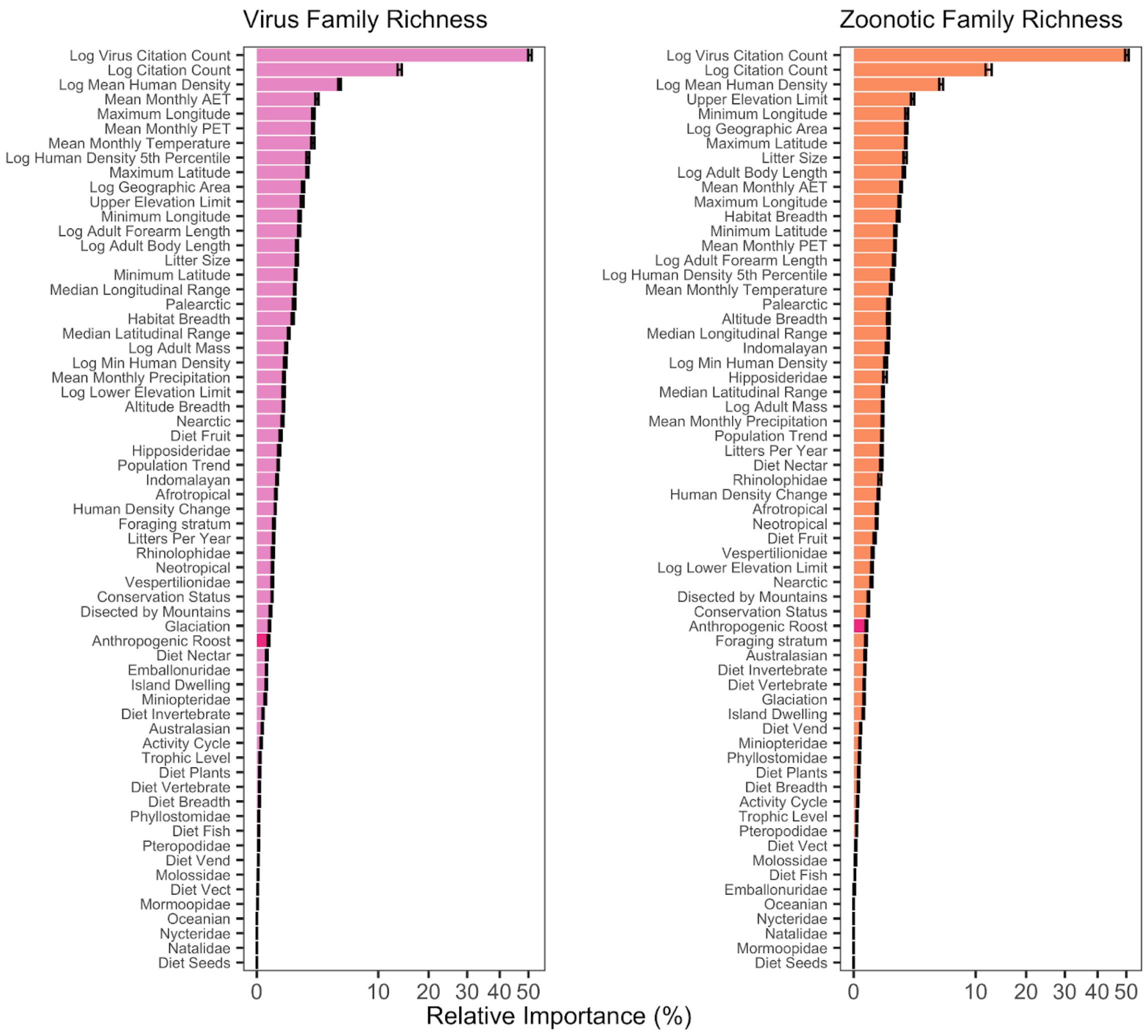

**Figure S4. Variable influence for predicting virus reservoir status and zoonotic reservoir status.** Bar height shows the importance of each predictor averaged across 100 unique test data splits. Error bars indicate the variation across models. Axes of relative importance are plotted on square root scale.

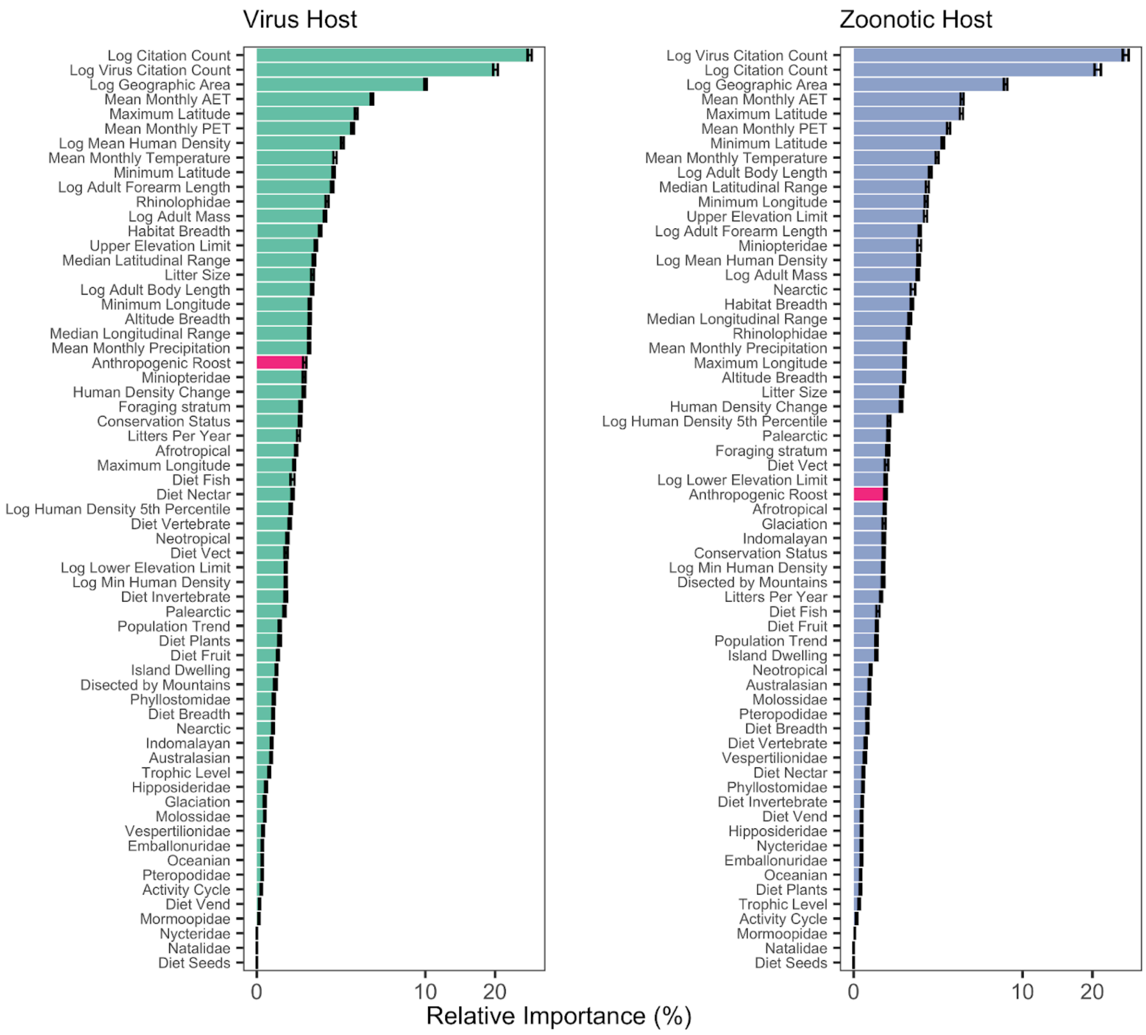

**Figure S5. Partial dependence plots of the top 10 variables for predicting virus family richness.** Pink lines and dots show the marginal effect for a single model run. Black lines and dots show the average across all 100 model runs. Histograms of numeric variables are shown in gray.

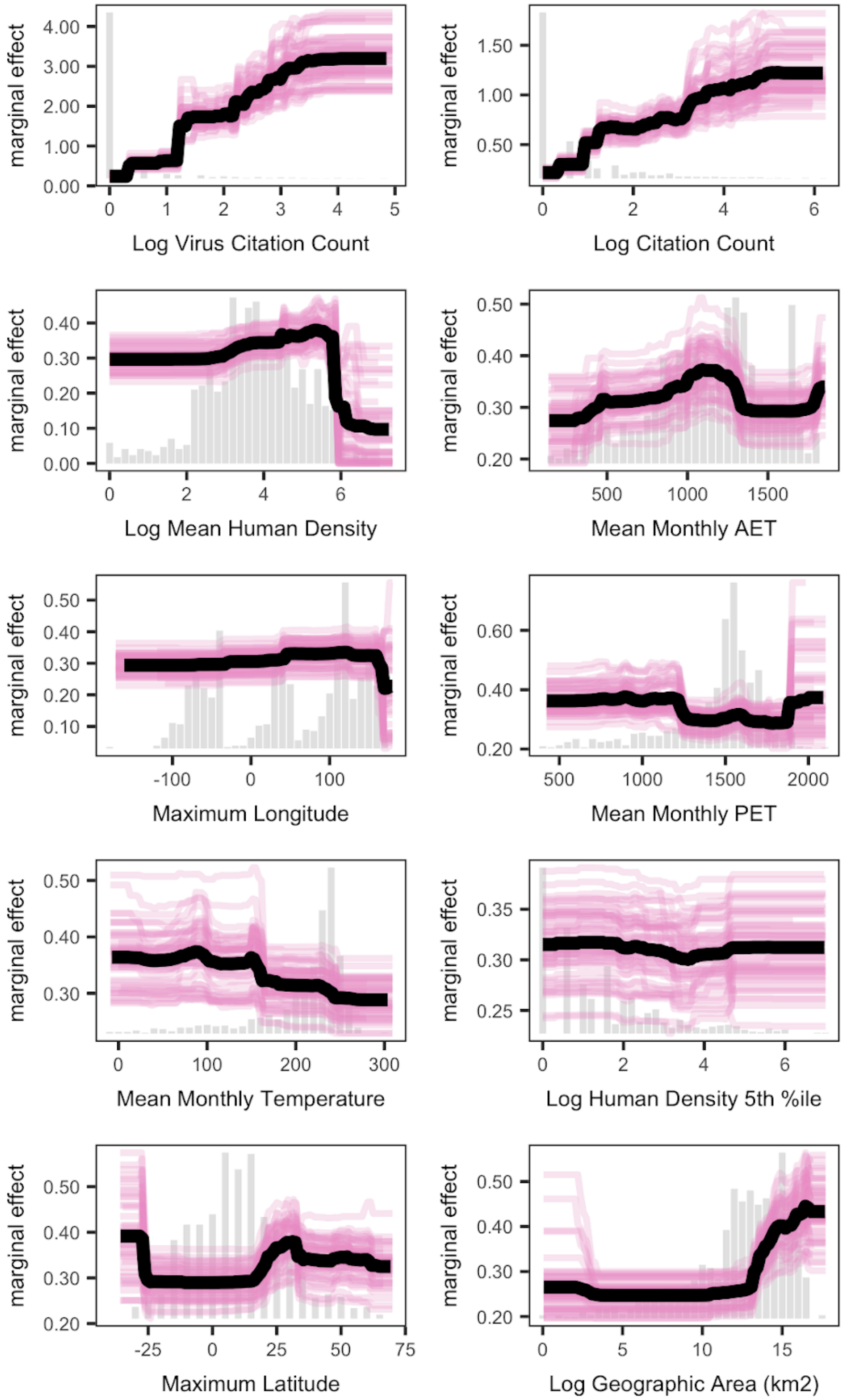

**Figure S6. Partial dependence plots of the top 10 variables for predicting zoonotic family richness.** Orange lines and dots show the marginal effects for individual model runs. Black lines and dots show the average across all 100 model runs. Histograms of numeric variables are shown in gray.

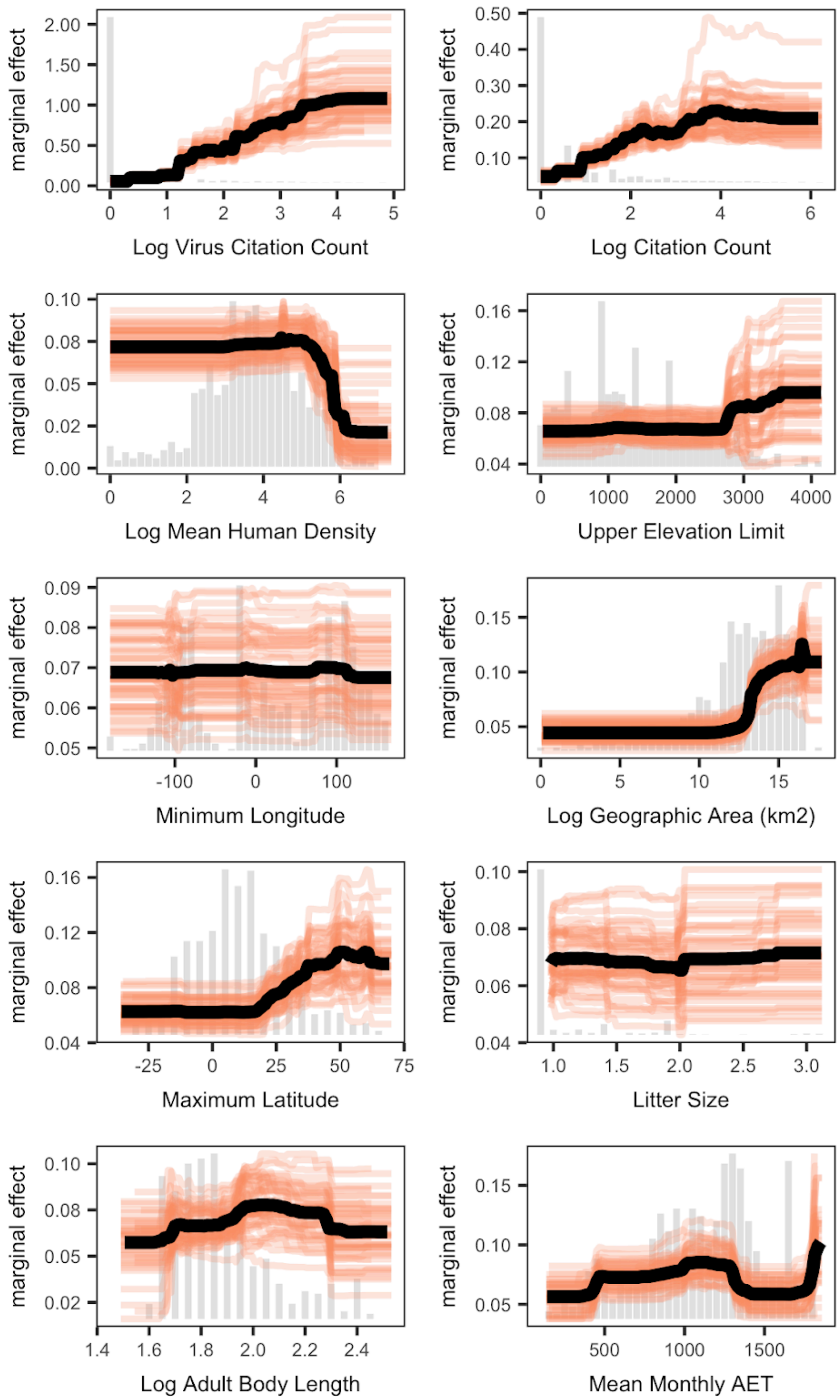

**Figure S7. Partial dependence plots of the top 10 variables for predicting virus host status.** Green lines and dots show the marginal effects for individual model runs. Black lines and dots show the average across all 100 model runs. Histograms of numeric variables are shown in gray.

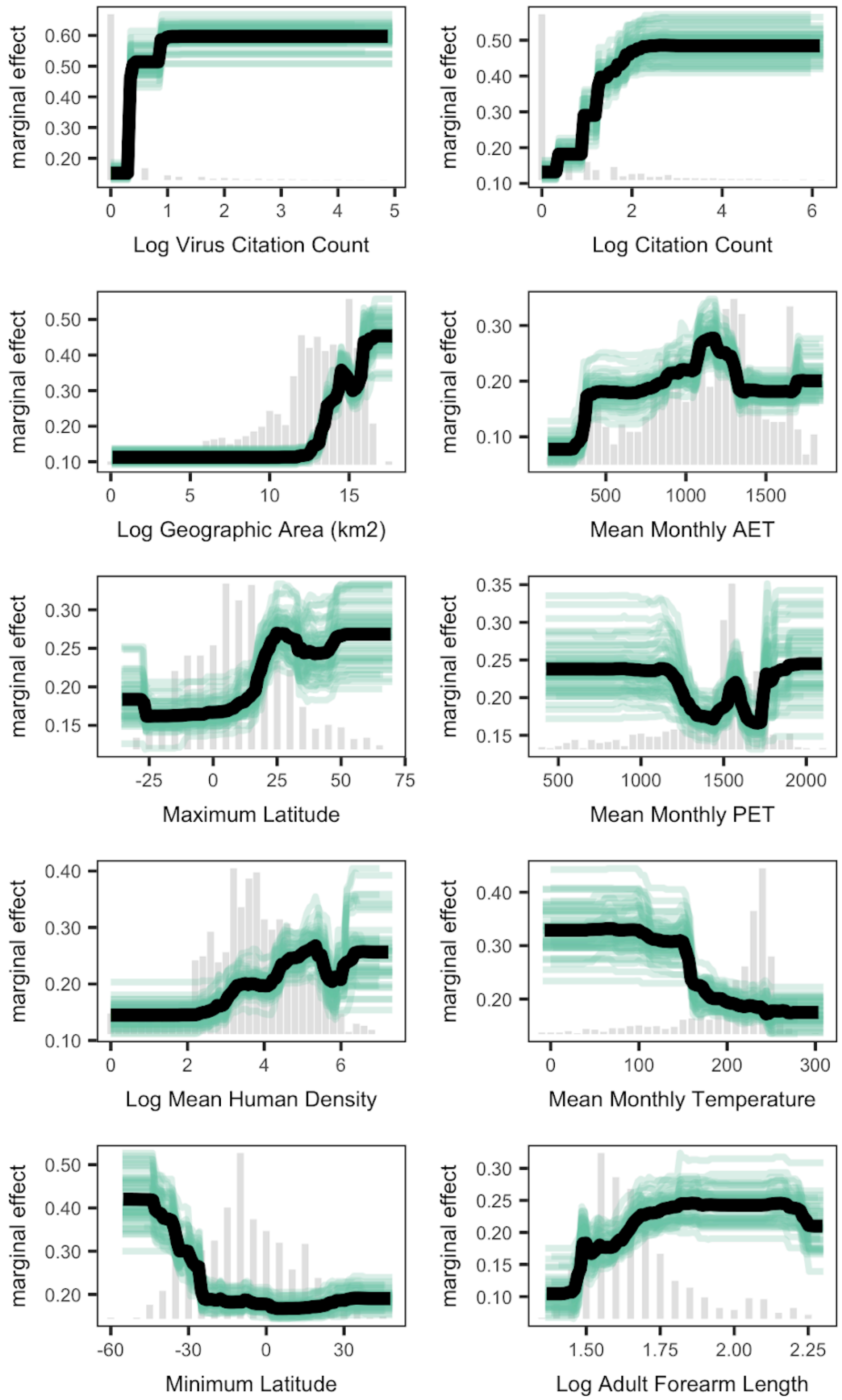

**Figure S8. Partial dependence plots of the top 10 variables for predicting zoonotic virus host status.** Blue lines and dots show the marginal effects for individual model runs. Black lines and dots show the average across all 100 model runs. Histograms of numeric variables are shown in gray.

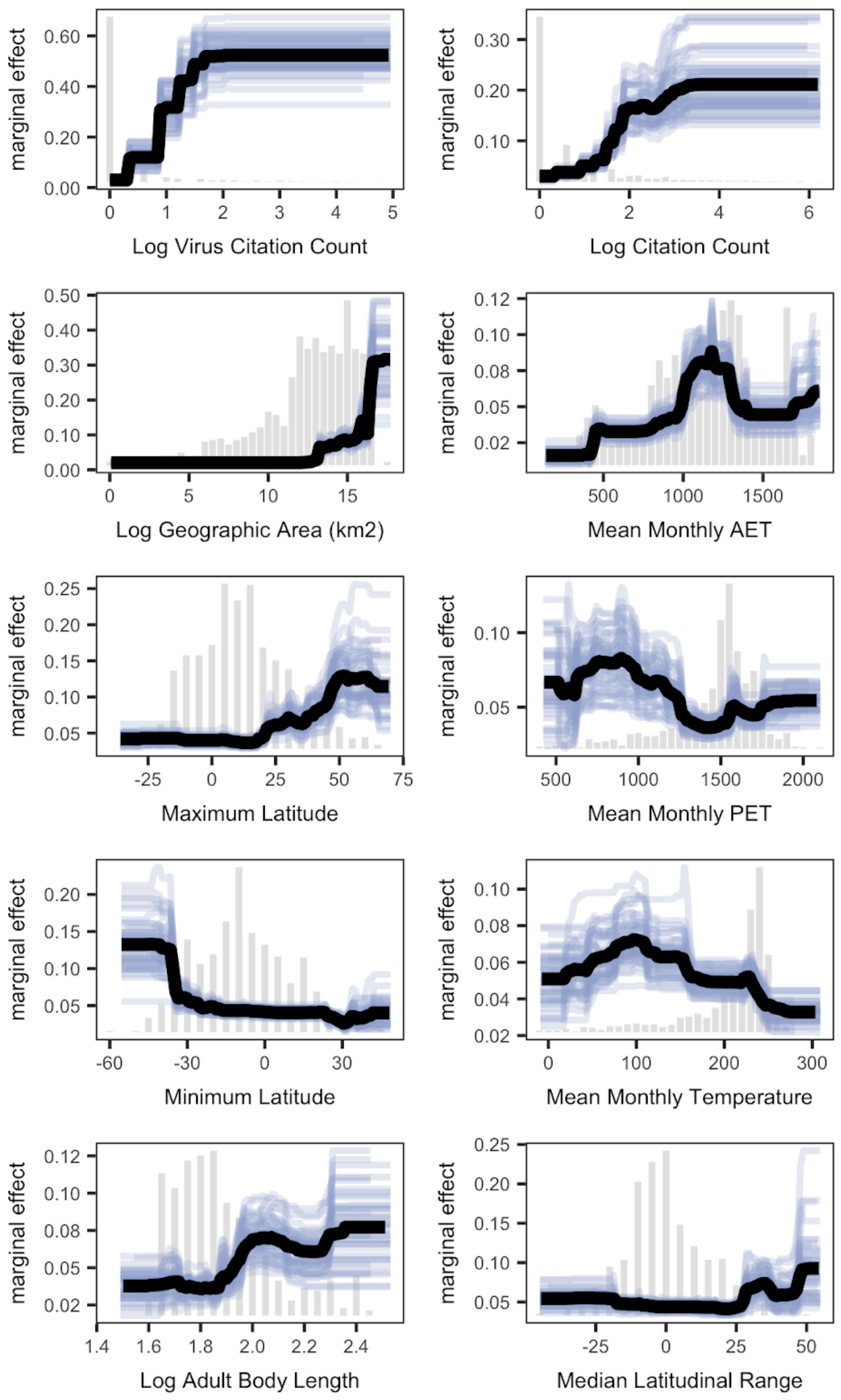

**Figure S9. Scatterplots of predictions from host models with and without anthropogenic roosting included as a predictor of virus family richness (A), zoonotic virus family richness (B), viral hosting ability (C) and zoonotic viral hosting ability (D).** Spearman rank correlation coefficients and p-values are displayed in the top left of both panels.

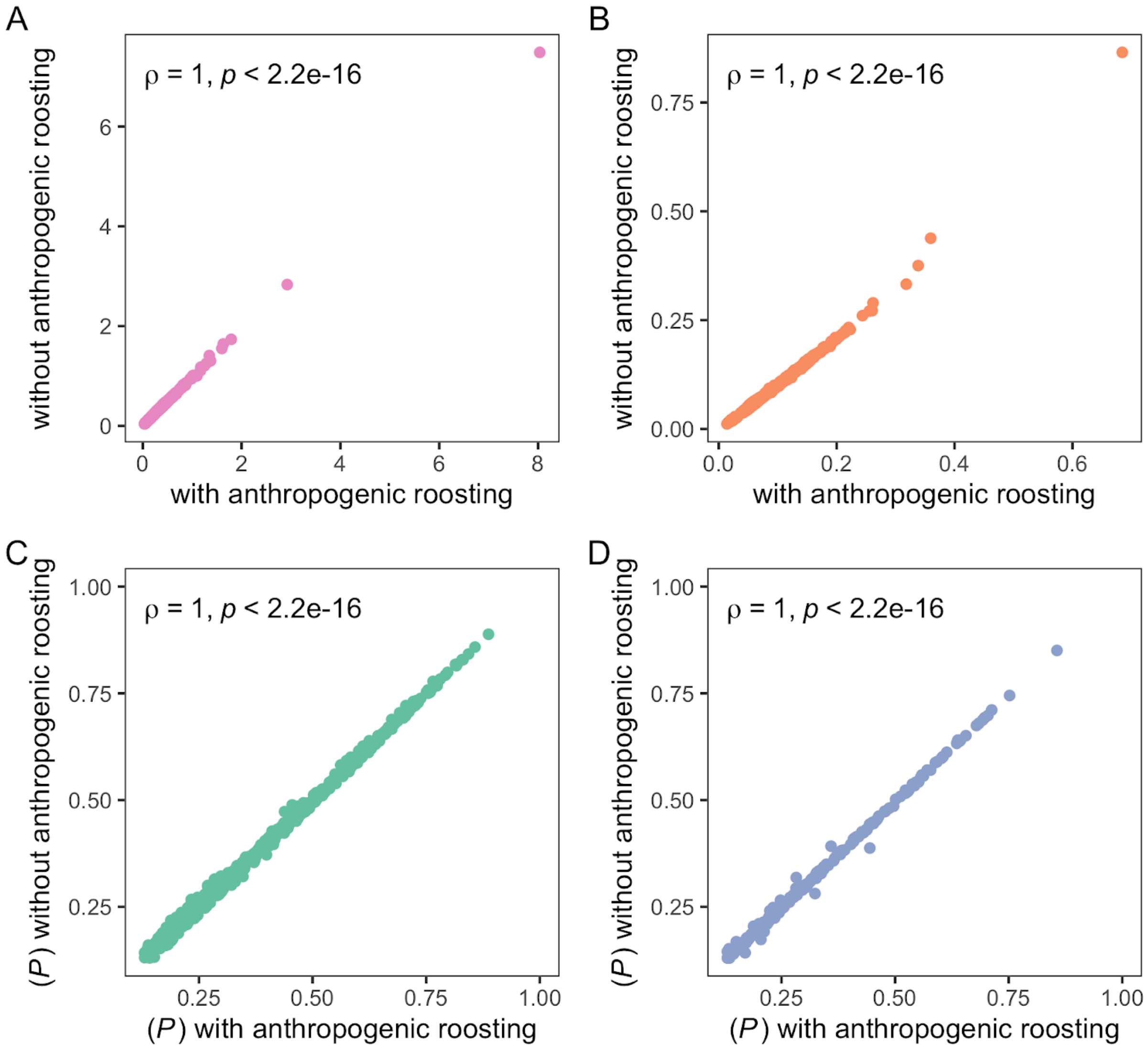

**Figure S10. Comparison of average relative importance between anthropogenic roosting and human population density metrics for virus hosting models.** Error bars represent the standard error of relative importance across the 50 model runs. Anthropogenic roosting is colored in red and human population density metrics are colored in grey.

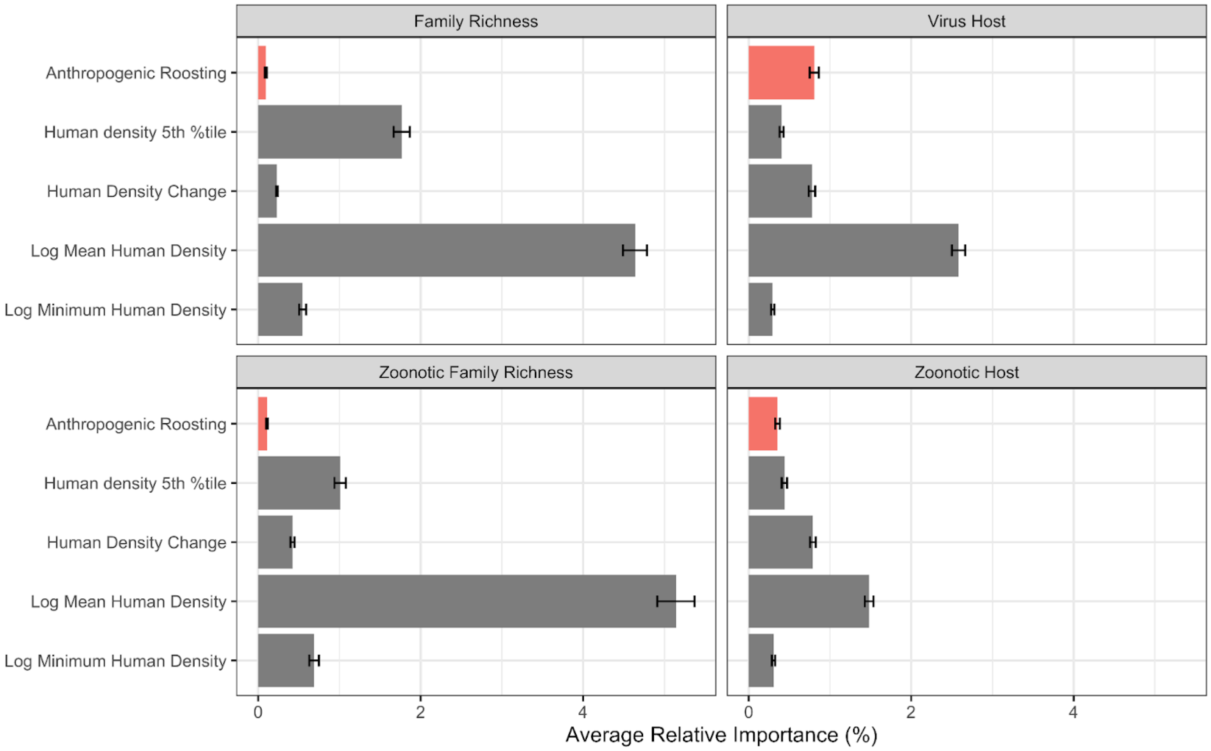

**Figure S11. Distribution of virus host diversity.** The first column shows the distribution of known anthropogenic (top left panel) and natural roosting (bottom left panel) hosts of viruses. The second column shows the distribution of undiscovered anthropogenic roosting (top right panel) and natural roosting (bottom right panel) hosts. Maps are based on the IUCN range list of mammal geographical ranges.

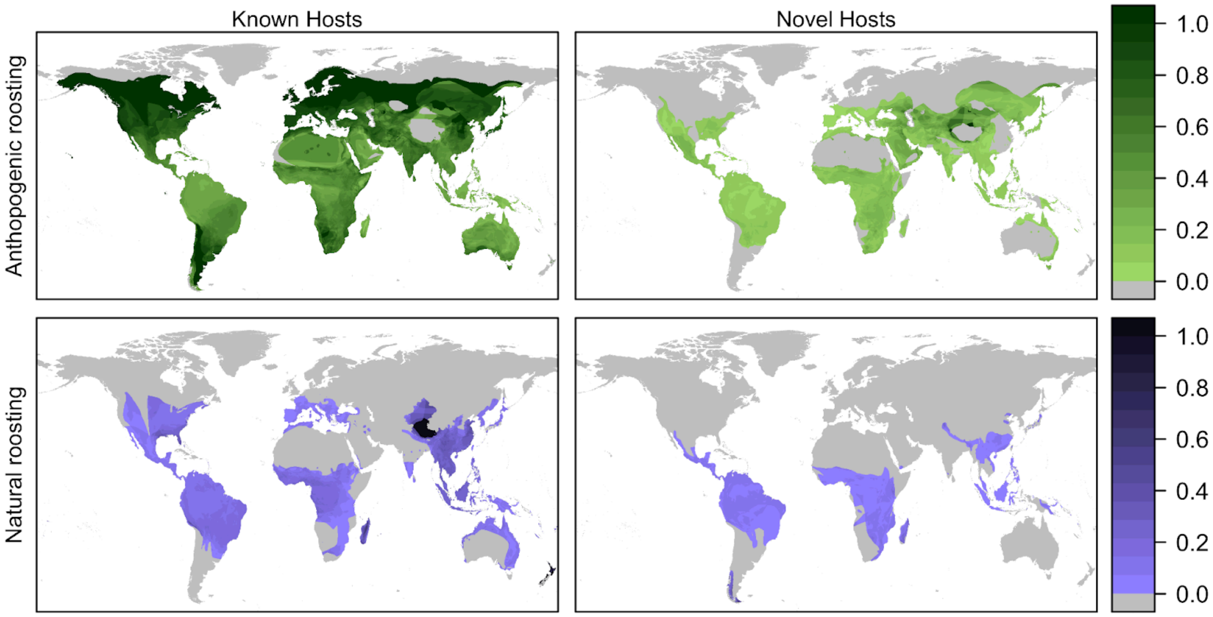

**Figure S12. Comparison of average relative importance between anthropogenic roosting and the 15 traits that characterize anthropogenic roosting across viral outcomes.** With the exception of the anthropogenic roosting variable, all other variables are in the order of most to least important for characterizing anthropogenic roosting as identified in Betke et al. 2024. The importance for Anthropogenic roosting is colored in red and the top metrics are colored in grey. Error bars represent the standard error of relative importance across the 50 model runs.

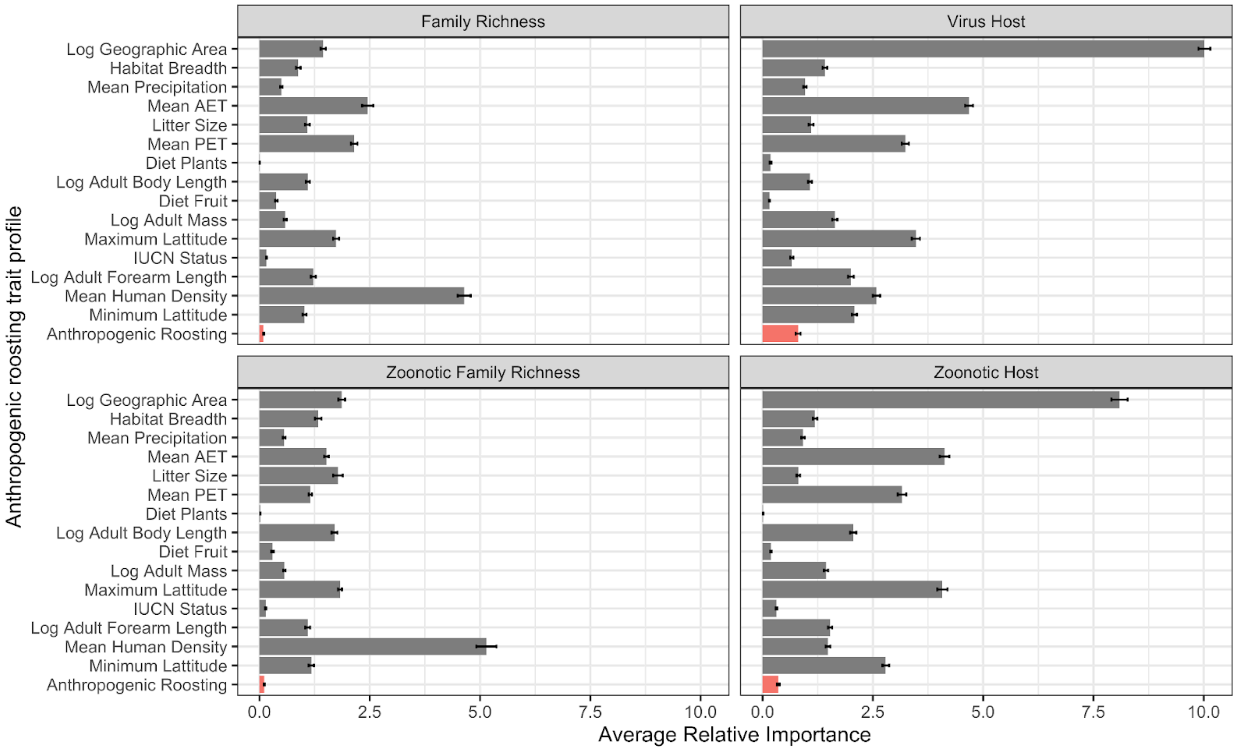

**Tables**

**Table S2. Summary of pairwise comparisons between all combinations of viral outcomes for feature ranking of anthropogenic roosting.** P-values were adjusted for multiple comparisons using the Benjamini–Hochberg correction.

| **contrast** | **estimate** | **SE** | **df** | **t.ratio** | **p.value** |
|---|---|---|---|---|---|
| virus richness - zoonotic richness | -0.70 | 1.2 | 196 | -0.582 | 0.5612 |
| virus richness - virus host | 16.32 | 1.2 | 196 | 13.569 | <.0001 |
| virus richness - zoonotic host | 7.04 | 1.2 | 196 | 5.853 | <.0001 |
| Zoonotic richness - virus host | 17.02 | 1.2 | 196 | 14.151 | <.0001 |
| zoonotic richness - zoonotic host | 7.74 | 1.2 | 196 | 6.435 | <.0001 |
| virus host - zoonotic host | -9.28 | 1.2 | 196 | -7.716 | <.0001 |
